# Oxygen carrying nanoemulsions and respiratory hyperoxia eliminate tumor hypoxia-induced suppression and improve cancer immunotherapy

**DOI:** 10.1101/2024.02.17.580835

**Authors:** Katarina Halpin-Veszeleiova, Michael Mallouh, Ashley Apro, Nuria Romero, Camille Bahr, Maureen Shin, Kelly Ward, Laura Rosenberg, Michail V. Sitkovsky, Bruce Spiess, Stephen M. Hatfield

## Abstract

Hypoxia-HIF-1α-driven immunosuppressive transcription and cAMP-elevating signaling through A2A-adenosine receptors (A2AR) represent a major tumor-protecting pathway that enables immune evasion. Recent promising clinical outcomes due to the blockade of the adenosine-generating enzyme CD73 and A2AR in patient’s refractory to all other therapies have confirmed the importance of targeting hypoxia-adenosinergic signaling. We report a novel and feasible approach to target the upstream stage of hypoxia-adenosinergic immunosuppression using an oxygen-carrying nanoemulsion (perfluorocarbon blood substitute). It is shown that oxygenation agent therapy i) eliminates tumor hypoxia, ii) improves efficacy of endogenously developed and adoptively transferred T cells, and thereby iii) promotes regression of tumors in different anatomical locations. We show that both T cells and NK cells avoid hypoxic tumor areas and that reversal of hypoxia by oxygenation agent therapy increases intratumoral infiltration of activated T cells and NK cells due to re-programming of the tumor microenvironment (TME). Thus, repurposing oxygenation agents in combination with supplemental oxygen may improve current cancer immunotherapies by preventing hypoxia-adenosinergic suppression, promoting immune cell infiltration and enhancing effector responses. These data also suggest that pretreating patients with oxygenation agent therapy may reprogram the TME from immune-suppressive to immune-permissive prior to adoptive cell therapy, or other forms of immunotherapy.

**Summary:** Oxygen delivering nanoemulsions and respiratory hyperoxia address limitations of blood vessel-mediated tumor oxygenation and promote anti-tumor immune responses to enhance immunotherapy.

## Introduction

The discovery of immune checkpoints motivated pre-clinical and clinical studies that have led to novel immunotherapeutic treatment modalities ^1-4^. While immune checkpoint blockade has afforded durable clinical responses, many patients relapse or fail to respond. In contrast to these immunological barriers, here we explored the targeting of hypoxia-adenosinergic biochemical negative regulators, which have been demonstrated in both preclinical studies and clinical observations to induce immune-suppression in the tumor microenvironment (TME).

Hypoxia-adenosinergic signaling emerged as a potent immunomodulating mechanism that governs the duration and direction of anti-pathogen and anti-tumor immune responses. It was established ^5-7^ and confirmed ^8-23^ that the major signaling components that drive this immune-suppressive axis are G_s_-protein coupled A2A adenosine receptors (A2AR) that trigger the accumulation of cAMP and subsequent activation of downstream PKA-mediated signaling events^24^. While this hypoxia-driven and extracellular adenosine-mediated biochemical pathway inhibits effector functions of many immune cell subtypes, it is most well documented in T cells, where transmembrane signaling from A2AR delivers an “off signal” to early and late events to TCR-signaling as well as phosphoCREB-mediated transcriptional changes in genes with cAMP response elements (CRE) (Fig. 1a) ^25^. In the context of the hypoxic TME, the anti-inflammatory hypoxia-HIF è extracellular adenosine è A2AR è cAMP è PKA signaling cascade leads to inhibition of the anti-tumor immune response via reduction of cytokine production, proliferation, cytolytic capacity, among other suppressive mechanisms ^5,7,26-33^.

**Fig. 1.**
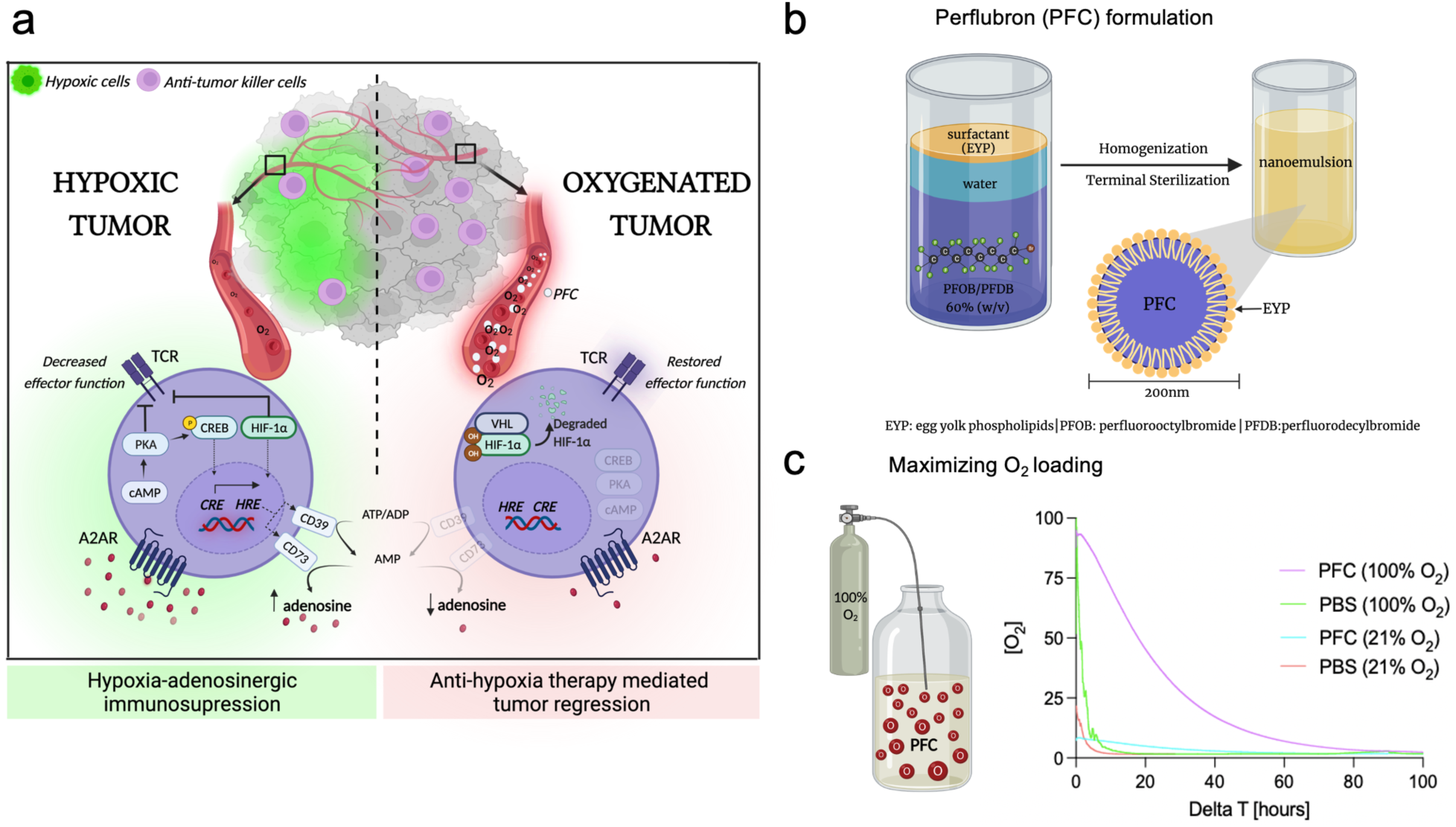
Exposure of perfluorocarbon nanoemulsion to 100% O_2_ prior to in vivo administration maximizes levels of dissolved oxygen. **(a)** Schematic representation of the rationale to use PFC to reverse tumor hypoxia and target hypoxia-adenosinergic signaling. The hypoxia/HIF1α-driven increase in adenosine generating ecto-enzymes CD39/CD73 leads to the accumulation of extracellular adenosine in the TME that triggers immunosuppressive A2AR → cAMP → PKA signaling. PFCs target the upstream stage of the hypoxia-adenosinergic axis of immunosuppression by increasing tumor oxygen tension and eliminating hypoxia/HIF1α and A2AR-mediated signaling. Tumor oxygenation weakens all-known stages of the hypoxia-adenosinergic signaling axis to prevent inhibition of anti-tumor cytotoxic killer cells and promote tumor regression ^28,29^. *(HRE; hypoxia response element, CRE; cAMP response element, PKA; Protein Kinase A; CREB; cAMP response element binding protein, HIF1α; hypoxia inducible factor-1α, VHL; Von Hippel-Lindau, PFC; perfluorocarbon, TCR; T cell receptor)*. **(b)** Illustration depicting Perflubron, a perfluorocarbon-based nanoemulsion consisting of 60% (w/v) perfluorooctylbromide (PFOB) and perfluorodecylbromide (PFDB) in water stabilized with egg yolk phospholipids (EYP). After homogenization nanoemulsion consists of particles which are less than 200 nm in size. The amount of dissolved oxygen (O_2_) in Perflubron is proportional to the partial pressure of O_2_ in the environment. **(c)** *Left*: Illustration depicting the strategy to maximize oxygen carrying capacity of Perflubron by saturation with 100% O_2_ for 7 minutes. Unsaturated samples were maintained at 21% O_2_. *Right:* One mL of unsaturated (21%O_2_) or saturated (100%O_2_) PFC or phosphate buffered saline (PBS) as control was placed into H35 HEPA Hypoxystation (Don-Whitley) maintained at 1% O_2_, 5% CO_2_ and 37°C. Dissolved oxygen concentration was measured continuously by Presens oxygen monitoring system and Presens in vitro oxygen probes.

The hypoxia-HIF-1α-driven and extracellular adenosine-mediated immunosuppressive axis is now established as a critical mechanism that protects cancerous tissues and drives resistance to cancer therapies. This was based on previous insights from genetic, biochemical, and pharmacological landmark studies ^5,7^ demonstrating that hypoxia-A2-adenosinergic immunosuppressive signaling functions as a general tissue-protecting mechanism to protect normal cells from excessive collateral damage by overactive anti-pathogen immune responses, but also protects cancerous cells. Hypoxia-adenosinergic immunosuppression was discovered in the context of inflammation, where damage to the endothelium and subsequent hypoxia increases the expression of adenosine-generating enzymes (e.g., CD39/CD73) ^10-13,34-38^ leading to the accumulation of extracellular adenosine that signals through cAMP-elevating A2A and A2B adenosine receptors. However, it was soon demonstrated that this fundamental, tissue-protecting mechanism is hijacked by tumor cells in the hypoxic, adenosine rich TME to suppress anti-tumor immune responses ^7^.

Of note, the hypoxia-adenosinergic biochemical barrier in the TME also promotes and strengthens immunological barriers by augmenting suppression by immune checkpoints. A2AR-mediated signaling has been shown to increase levels of immunological negative regulators including LAG-3, IL-10, and T regulatory cells (T regs), among others ^15,39^. Additional lines of evidence also suggest that hypoxia/HIF-1α signaling promotes expression of immunological barriers such as CTLA-4, LAG-3, PD-1 and TIM-3 ^28,40^. Crosstalk between hypoxia/HIF-1α and A2-adenosinergic signaling has been further confirmed in studies of cAMP-dependent PKA signaling, where phosphorylation of PKA was shown to recruit the transcriptional activity of HIF-1α ^41^. Furthermore, the adenosine-generating ectoenzymes CD39/CD73 are regulated by HIF-1α and have been shown to promote tumor growth, invasiveness and metastatic potential resulting in poor clinical outcomes ^8,9,16-20,28,29,35,36,42^.

The discovery of this tumor-protecting biochemical pathway advanced our understanding of pharmacological approaches to block the early stages (hypoxia/HIF1) and late stages (adenosine-A2AR-cAMP) of this signaling axis. Preclinical evidence for this approach employing anti-hypoxia and anti-A2AR strategies has led to the justification of multiple clinical trials using such drugs against cancers that are refractory to all other therapies. Therefore, there is an acute need to improve outcomes of patients who are unresponsive to treatments such as immune checkpoint blockade since these tumors are still protected by the hypoxia-adenosinergic immunosuppressive barrier.

Here, we provide pre-clinical evidence for the improvement of cancer immunotherapies by targeting hypoxia, the upstream stage of the hypoxia-adenosinergic axis, using an oxygenation agent, perfluorocarbon (PFC), in combination with respiratory hyperoxia (hereafter referred to as ‘oxygenation agent therapy’) ^43^. PFCs may address limitations of respiratory hyperoxia which may be ineffective in increasing oxygen levels in tumor areas that are not reached by healthy blood vessels, have significant vascular damage, or are in anatomical areas that may not be as oxygen-privileged as the lungs. PFCs overcome these impediments since they are significantly smaller in size compared to erythrocytes and oxygen is physically dissolved in the PFC nanoemulsion, rather than chemically bound as in hemoglobin ^44-47^. We suggest here the conceptually novel and immunologically-motivated use of blood substitutes (e.g. oxygenation agents) to enhance oxygen delivery to the TME, thereby weakening hypoxia-adenosinergic immunosuppression and improving cancer immunotherapies ^43^. This approach differs from previous strategies utilizing PFCs to enhance the therapeutic benefits of radiotherapy, phototherapy, and chemotherapy ^45,48,49^.

We hypothesized that hypoxic elimination following treatment with oxygenation agents will improve immunotherapeutic protocols since our previous work demonstrated that increasing oxygen concentration in the TME interrupts the downstream CD73/CD39 → adenosine → A2AR→ cAMP immunosuppressive axis (Fig. 1a) ^28,29^. These studies were the first to demonstrate the reversal of hypoxia by respiratory hyperoxia (60% O_2_) could reprogram the immunosuppressive metabolome, proteome and cytokinome profile of the hostile TME and improve anti-tumor responses ^28,29^. In this report, we present our findings on the synergy between two anti-hypoxia treatments – respiratory hyperoxia and oxygen carrying nanoemulsions.

These studies may resurrect an entire industry of blood substitutes that were developed previously for other indications including stroke, military applications, organ preservation and as blood substitutes during the AIDS epidemic. Our unique motivation to use oxygen-carrying nanoemulsions to eliminate tumor hypoxia as the upstream stage of the hypoxia-adenosinergic immunosuppressive axis provides rationale to repurpose these drugs to improve cancer immunotherapy. Importantly, perfluorocarbon nanoemulsions have previously been tested in over 3,000 patients and shown to be safe for other indications. Support for such an approach is also provided by pre-clinical and clinical studies by Curran’s group demonstrating improved therapeutic efficacy utilizing a related strategy with a hypoxia-targeting prodrug, Evofosfamide^23,50^.

## Results

### Optimizing oxygen delivery by perfluorocarbon nanoemulsions to target the upstream stage of hypoxia-adenosinergic signaling axis

Low levels of O_2_ (PO_2_ < 10mmHg ∼ 1.4% O_2_) in the TME are associated with significantly lower disease-free survival in cancer patients when compared to patients with higher mean intratumoral PaO_2_ ^51-53^. We have proposed that the hypoxic TME is a major obstacle preventing success of cancer immunotherapies (Fig. 1a), including adoptive cell therapy (ACT) and immune checkpoint blockade (ICB). While recent attention has focused on immunological barriers (ICB) in the TME, we have provided justification for the targeting of biochemical barriers (e.g. hypoxia-adenosinergic suppression) that hinder infiltration, proliferation and effector functions of anti-tumor immune cells ^23,24,28,29^. Our previous work has shown that hypoxic and adenosine-rich tumors are protected from attack by anti-tumor immune cells ^28,29,33^.

To address limitations of oxygen delivery via erythrocytes, we investigated the use of the perfluorocarbon (PFC)-based oxygenation agent, Perflubron, as an approach to target tumor hypoxia, the upstream stage of the hypoxia-adenosinergic immunosuppressive signaling axis. We hypothesized that the small size (<0.2 µm) of these particles will allow better penetration in narrow microvascular channels of solid tumors ^54^. Fig. 1b depicts the composition of Perflubron (PFC), a mixture of 60% (w/v) perfluorooctylbromide (PFOB) and perfluorodecylbromide (PFDB) in water stabilized with egg yolk phospholipids (EYP) ^47^. Here, we exploited the oxygen solubility of PFC which is directly proportional to the oxygen partial pressure in the surrounding environment (Henry’s Law) ^54^. Perflubron “off-the-shelf” has a concentration of dissolved O_2_ similar to atmospheric air (∼21% O_2_). We hypothesized that oxygenating Perflubron by exposure to 100% O_2_ would maximize the amount of dissolved oxygen in the PFC nanoemulsion prior to in vivo administration, thereby increasing oxygen delivering capacity (Fig. 1c, *left*).

As shown in Fig. 1c, we exposed Perflubron to 100% O_2_ for 7 minutes and compared the oxygen dissolving and release capacity of oxygenated Perflubron to the “off-the-shelf” preparation of Perflubron. To mimic the hypoxic TME, these in vitro studies were conducted in an H35 Hypoxystation (Don-Whitley) in 1% O_2_/5% CO_2_ using Presens oxygen monitoring system to measure levels of dissolved oxygen. Fig. 1c (*right*) demonstrates that the oxygen carrying capacity of oxygenated Perflubron was increased and the oxygen release profile was prolonged compared to control, non-oxygenated Perflubron. These in vitro studies suggest a potential benefit of exposing Perflubron to high oxygen prior to administration to enhance oxygen carrying capacity, which might also be achieved in vivo by combining with respiratory hyperoxia. The data from Fig. 1 provided a methodological rationale for i) oxygenating Perflubron prior to in vivo administration for maximized initial O_2_-loading and ii) combination therapy with respiratory hyperoxia during Perflubron administration (oxygenation agent therapy) to achieve higher fractions of inspired oxygen (F_i_O_2_) and maximize the amount of dissolved O_2_ in Perflubron after in vivo administration.

### Therapeutic targeting of hypoxia-governed immunosuppression via oxygenation agent therapy

To alleviate the inhibition of anti-tumor immunity in hypoxic tumors, we tested whether oxygenation agent therapy could reverse tumor hypoxia and reprogram the TME ^28^. For these studies, mice with established MCA205 fibrosarcoma intradermal tumors were given daily administration of Perflubron for 72 h with and without respiratory hyperoxia (60% O_2_). In Fig. 2a, representative immunofluorescent images (left) and quantification (right) of hypoxyprobe staining of tumor sections demonstrate that oxygenation agent therapy significantly reversed hypoxia in intradermal tumors. While a decrease in hypoxic regions was observed in mice receiving Perflubron alone, the near complete elimination of hypoxia was seen in mice treated with Perflubron in combination with respiratory hyperoxia (60% O_2_) (Fig. 2a and Supplemental Fig. 1a). These data were confirmed by flow cytometry assays demonstrating that the percentage and mean fluorescent intensity (MFI) of hypoxic tumor cells was significantly decreased in mice treated with Perflubron in combination with respiratory hyperoxia (Fig. 2b, Supplemental Fig. 1b). We then asked whether we could measure the increased oxygen levels in the tumor following Perflubron administration in real time. To this end, we employed Presens oxygen monitoring system to measure oxygen levels in anesthetized mice bearing intradermal tumors. After locating hypoxic tumor areas (< 1% O_2_) using an in vivo Presens oxygen probe, we infused Perflubron and continuously monitored oxygen tension. Twelve minutes after administration of Perflubron, the intratumoral oxygen levels increased to ∼15% O_2_ (pO_2_ ∼107 mmHg) and was sustained for the duration of the assay (∼250 minutes). Fig. 2c shows the real time measurements of the increase in oxygen concentration over the 4 h time period.

**Fig. 2.**
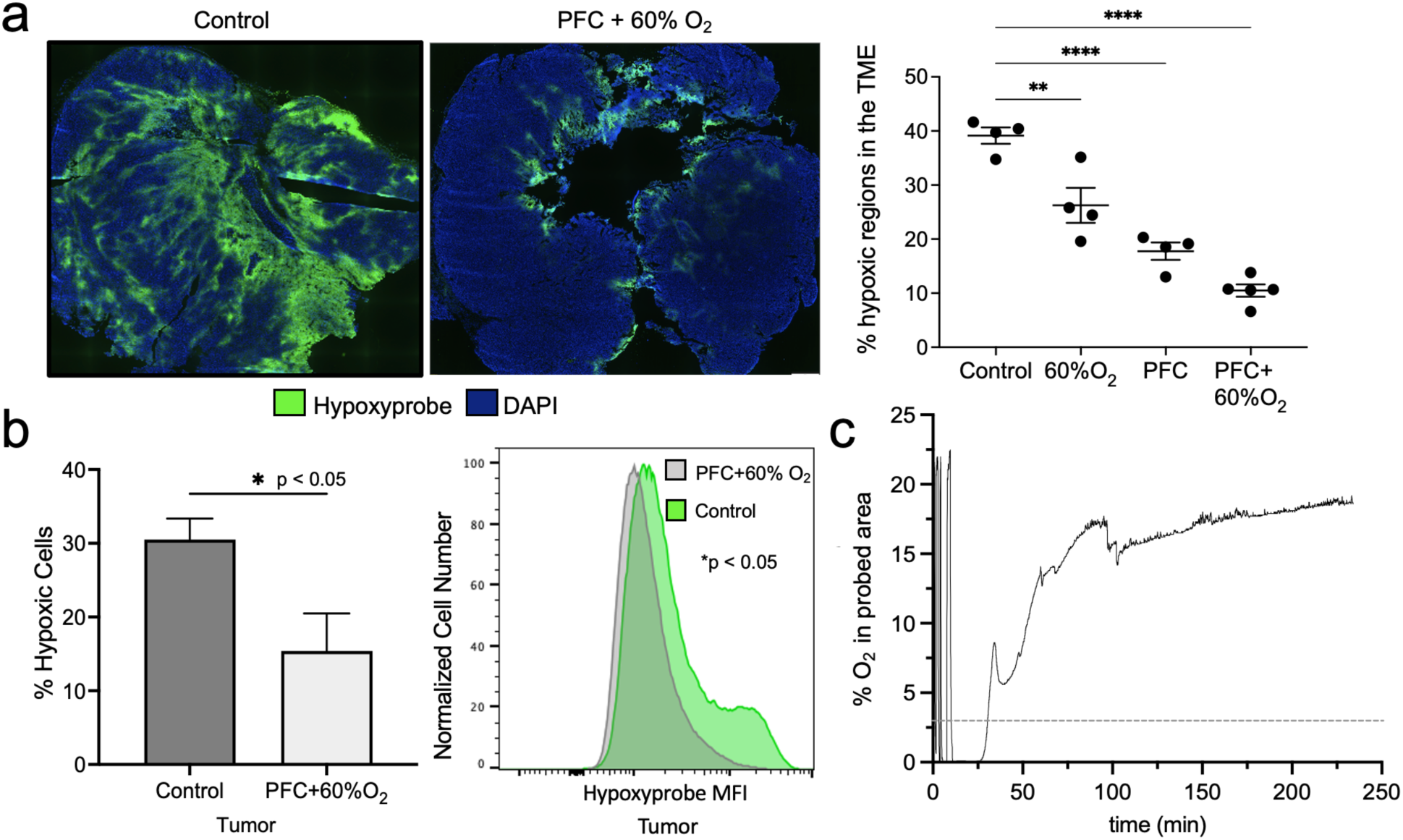
Oxygenation agent therapy eliminates intratumoral hypoxia. Mice with 8-day established MCA205 intradermal tumors were treated with or without i.v. administration of 15 ml/kg oxygenated Perflubron (PFC) for 72 h with continuous breathing of 60% O_2_ (respiratory hyperoxia) or 21% O_2_ (normoxia). Intra-tumoral hypoxia was measured by immunofluorescence (a) and (b) flow cytometry using the hypoxia marker Hypoxyprobe-1^™^ injected i.p. (80mg/kg) one hour prior to sacrifice. **(a)** *Left*: Representative fluorescence images of hypoxic levels (green) in the isolated tumors from control versus mice treated with PFC + 60% O_2_. *Right*: quantification of intratumoral hypoxia using CellSens software (Olympus) in all experimental groups. *P*-values are calculated using one-way ANOVA with Tukey post-test for multiple comparisons (*P<0.05, **P<0.005, ***P<0.0005, ****P<0.0001. Data presented as mean ± SEM, n ≥ 4). **(b)** *Left:* Tumors isolated from mice treated with oxygenation agent therapy (white bars) had significantly less hypoxic cells compared to control (gray bars) as measured by flow cytometric analysis of Hypoxypobe staining. *Right:* Increased mean fluorescent intensity (MFI) of hypoxyprobe staining on cells isolated from tumors from control mice (green) and mice treated with oxygenation agent therapy (gray). *P*-values are calculated using one-way ANOVA with Tukey post-test for multiple comparisons (*P<0.05, n ≥ 4). **(c)** Real-time measurement of increased intratumoral oxygen concentration in intradermal tumors after Perflubron infusion measured by Presens oxygen monitoring system and Presens in vivo probes. After stabilization of the oxygen probe in the hypoxic tumor region (0.02% O_2_) in an anesthetized mouse, Perflubron was infused i.v. (15 mL/kg) at minute 12. Oxygen concentration was recorded up to 229.5 minutes. The oxygen concentration curve at the probed tumor area is shown as a function of time before and after Perflubron administration (arrow indicates Perflubron administration and dotted line indicates oxygen concentration where HIF-1α is considered destabilized). Demonstration of increased intratumoral oxygen tension by Perflubron administration was repeated in three independent experiments.

Since our goal is to decrease the hypoxia-HIF-1α-mediated triggering of immunosuppressive downstream events, it is important to emphasize that 10-15 mmHg (1.5%-2% O_2_) is considered 50% inhibitory concentration of the catalytic subunit of HIF-1α ^53,55^. Therefore, Fig. 1 and 2 represent mechanistic biochemical justification for using our approach in vivo since, according to published reports ^56^, at the partial pressure of oxygen achieved by oxygenation agent therapy (PO_2_ 107mmHg), HIF-1α is destabilized ^56^. According to clinical data, such increase in oxygen concentration has dramatic consequences in the responses and outcomes of cancer patients. Of note, previous work by Dewhirst and colleagues showed that patients with intratumoral PaO_2_ >11 mmHg have over 50% higher chance of disease-free survival^52^.

### Oxygenation agent therapy reprograms the TME to promote anti-tumor immune cell infiltration

Since we proposed that tumor hypoxia is a powerful immune-suppressive barrier, we then examined the infiltration of anti-tumor immune cells into normoxic versus hypoxic regions of tumors. Data from Fig. 3a extend previous observations by demonstrating that both anti-tumor CD8 T cells and NK cells avoid hypoxic regions of intradermal tumors as measured by the biochemical hypoxia marker, Hypoxyprobe ^23,28^. Significantly fewer CD8 T cells (Fig. 3a) and NK cells (Fig. 3b) were observed in hypoxic tumor areas (<1% O_2_) in all of the intradermal tumors assayed.

**Figure 3.**
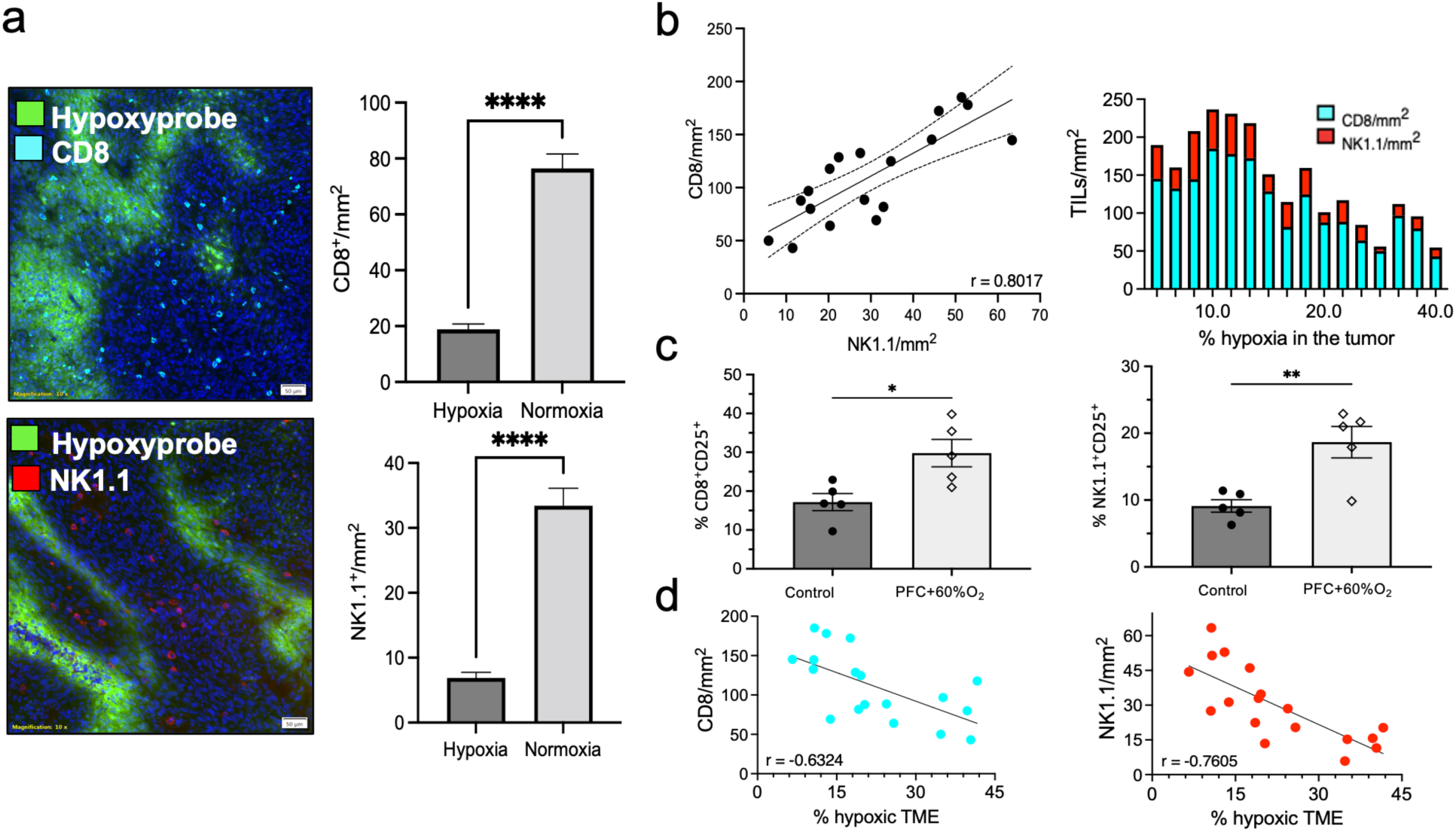
Oxygenation agent therapy reprograms the tumor microenvironment toward immune-permission. **(a)** Anti-tumor T cells and NK cells avoid hypoxic regions of solid tumors. MCA205 intradermal tumors were isolated and 5 µM frozen tissue sections were prepared. Fixed sections were fluorescently labeled with anti-Hypoxyprobe, anti-CD8 and anti-NK1.1 antibodies to determine infiltration of immune cells into the hypoxic (green) or non-hypoxic regions (dark blue, DAPI). *Left panels:* representative fluorescence images demonstrate (A) CD8+ T cells (top, light blue) and (B) NK cells (bottom, red) avoid hypoxic regions (green). *Right panels:* quantification of CD8 T cell (top) and NK cell (bottom) infiltration into hypoxic vs. non-hypoxic areas. A total of 60 different regions from the TME across 15 tumors were analyzed using CellSens software (Olympus) and results are shown in the bar graphs (right) (****P<0.0001, data presented as mean ± SEM. The scale bar represents 50 µm,10x magnification). **(b-d)** C57BL/6 mice with 8-day established MCA205 intradermal tumors received daily administration (i.v., 15 ml/kg) of oxygenated Perflubron for 72 h while breathing 60% O_2_ (respiratory hyperoxia) or 21% O_2_ (normoxia). Mice were injected intraperitoneally with Hypoxyprobe-1^™^ (80mg/kg) one hour before sacrifice to determine levels of intra-tumoral hypoxia. After sacrifice, tumors were isolated divided for analysis via immunofluorescence or flow cytometry. **(b)** *left:* Quantitative analysis using CellSens software (Olympus) of 5 μM thick frozen tumor sections correlating the number of infiltrating NK cells and CD8 cells per tumor. Each dot represents one tumor (n = 18). Dotted line represents 95% confidence interval. *right:* Bar graph depicting the relationship between NK cells (red) and CD8 cells (blue) and levels of intratumoral hypoxia. Each bar represents one tumor and corresponding numbers for infiltration and hypoxic regions (n = 18). **(c)** Quantitation of the numbers CD8+CD25+ (left) and NK+CD25+ (right) cells from tumors from mice treated with oxygenation agent therapy versus control. *P*-values are calculated using one-way ANOVA with Tukey post test for multiple comparisons (*P<0.05. Data are presented as mean ± SEM, n = 5). **(d)** Pearson’s *c*orrelation analyses between the percentage of hypoxic regions in the TME and the number of infiltrating CD8 T cells (left) or NK cells (right). Each dot represents one tumor (*n* = 18).

The absence of CD8 T cells and NK cells in hypoxic regions led us to investigate whether reprogramming the TME by reversing tumor hypoxia (Fig. 2) ^29^ ^28^ might affect recruitment of endogenous anti-tumor immune cells. Using fluorescent microscopy, we quantified the numbers of CD8 T cells, NK cells and hypoxia from intradermal tumors. Mice were treated with daily administration of Perflubron and respiratory hyperoxia (60% O_2_) for 72 h or kept at atmospheric oxygen levels as a control. We first evaluated the relationship between infiltrating CD8 T cells and NK cells regardless of hypoxic levels in the TME. Figure 3b (left) shows a strong positive correlation between the number of infiltrating T cells and NK cells into tumors (r = 0.8), supporting previous observations that NK cells may recruit and orchestrate activities of CD8 T cells ^28^. We then examined the number of T cell and NK cell infiltrates in the context of tumor hypoxia. Fig. 3b (right) demonstrates that tumors with a lower percentage of hypoxic regions had higher levels of infiltrating T cells and NK cells. These observations are in agreement with findings from Fig. 3a. Taken together, these data suggest that the mechanisms that attract or inhibit T cell or NK cell infiltration are strongly affected by tumor hypoxia. This is supported by our previous findings that the reversal of hypoxia significantly increased the expression of proinflammatory cytokines and chemokines in the TME ^28^.

Next, we asked whether reprogramming of the TME by oxygenation agent therapy would increase the recruitment and activation of anti-tumor immune cells. Fig. 3c and Supplemental Fig. 2 show that oxygenation agent therapy resulted in significantly increased numbers of intratumoral CD8^+^CD25^+^ (*left*) and NK1.1^+^CD25^+^ (*right*) cells. We then examined the correlation between hypoxia and CD8 T cells or NK cells. Fig. 3d demonstrates a strong negative correlation between CD8 T cell infiltration and hypoxia (r = -0.6324, left), and an even stronger negative correlation between NK cells and hypoxic levels (r = -0.7605, right). These data suggest that NK cells may be more sensitive to immunosuppressive hypoxic signaling. This supports previous demonstrations of the high susceptibility of NK cells to adenosine-A2AR-mediated suppression, which is prominent in hypoxic conditions ^28,57,58^.

### Oxygenation agent therapy induces tumor regression and improves survival

After demonstrating that oxygenation agent therapy eliminates tumor hypoxia and increases the infiltration of tumor reactive lymphocytes, we tested whether this approach is capable of inducing tumor regression. Fig. 4 a,b show that while Perflubron alone and respiratory hyperoxia (60% O_2_) alone resulted in a slight tumor growth delay, oxygenation agent therapy (3 Perflubron injections/week + 60% O_2_) promoted significant regression of 9-day established intradermal MCA205 fibrosarcoma tumors compared to control mice. This potent tumor regression induced by oxygenation agent therapy was also confirmed in the CT26 colon carcinoma model in Fig. 4d,e. These data extend previous findings of the immunological anti-tumor effects of respiratory hyperoxia in lung tumor models that are much more readily oxygenated by breathing 60% O_2_ alone ^28,29^. Interestingly, we observed striking ulceration and necrosis in tumors from mice treated with oxygenation agent therapy, which may indicate morphological signs of tumor destruction by anti-tumor immune cells (Fig. 4e, insert). This phenomenon was not observed in tumors from other groups. Additional mechanistic assays confirmed that these tumor-bearing mice treated with oxygenation agent therapy exhibited decreased intratumoral hypoxia at the study end point (Supplemental Fig. 3). Figure 4c,f demonstrate that oxygenation agent therapy also significantly improved the survival of intradermal tumor-bearing mice compared to untreated control mice in two separate tumor models (CT26 and MCA205). Taken together, these data suggest that oxygenation agent therapy eliminates tumor hypoxia, promotes infiltration of activated anti-tumor immune cells and induces significant tumor regression, particularly in anatomical locations that are not as oxygen privileged as the lung.

**Figure 4.**
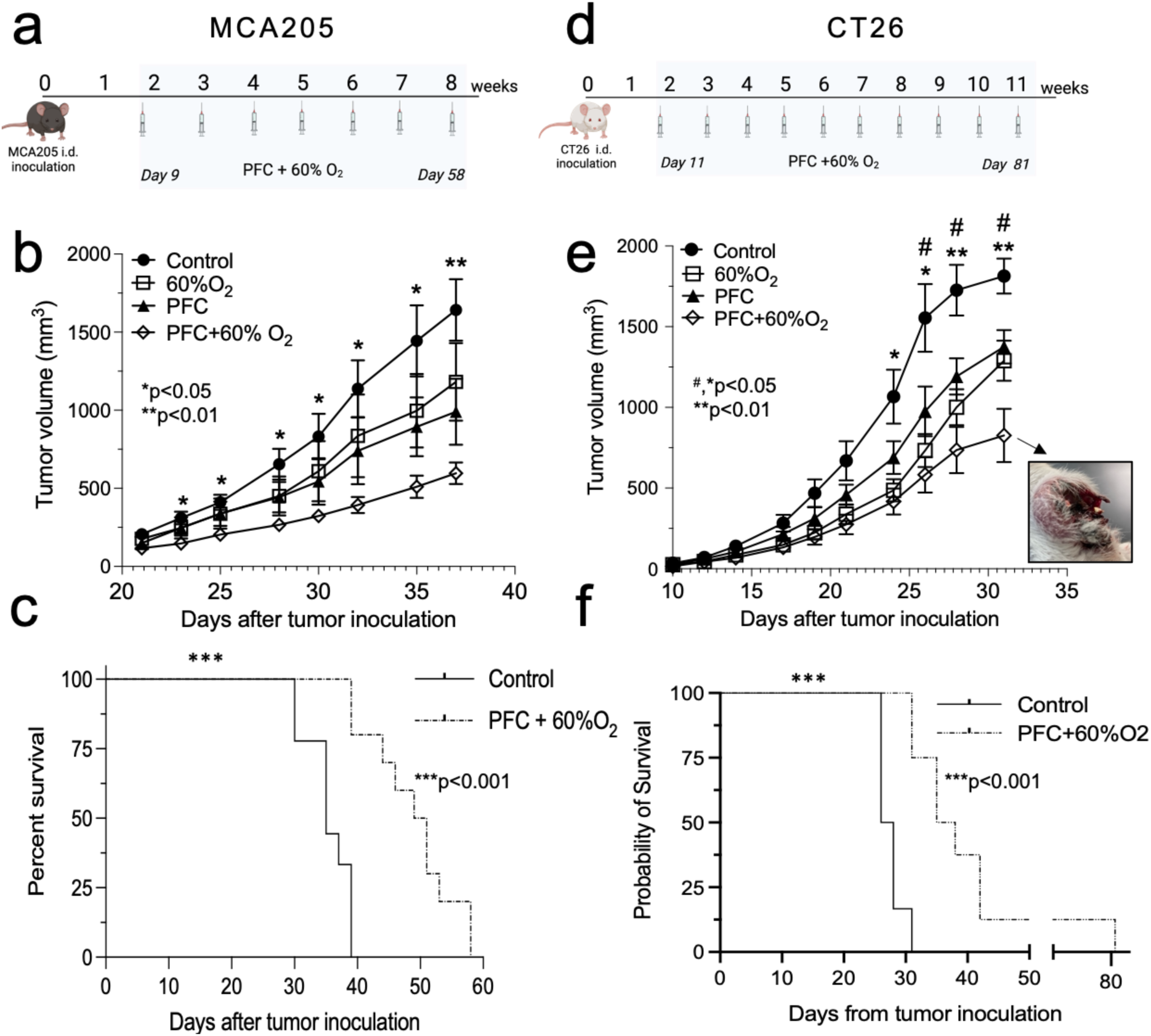
Oxygenation agent therapy induces tumor regression and improves survival. C57BL/6 and BALB/c mice were injected intradermally with MCA205 fibrosarcoma or CT26 colon carcinoma and treated with or without Perflubron (PFC) and respiratory hyperoxia (60% O_2_) **(a)** Schematic illustration of the experimental design and treatment regimen in MCA205 intradermal model. C57BL6 mice with 9-day established MCA205 fibrosarcoma intradermal tumors were housed in 60% O_2_ chambers or maintained at 21% O_2_ (normoxia) and treated with or without oxygenated Perflubron (15 mL/kg) 3 times per week until study completion. **(b)** Oxygenation agent therapy (PFC + 60% O_2_) induces the strongest tumor regression compared to control or either treatment alone. Tumors were measured three times per week using Vernier calipers and tumor volume was calculated by the following formula: 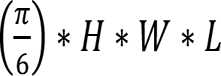. Tumor sizes were assessed by two-way analysis of variance (ANOVA) with multiple comparisons (*P<0.05, **P<0.005. Data presented as mean ± SEM, *n* ≥ 7). **(c)** Survival curves for MCA205 tumor-bearing mice treated with oxygenation agent therapy (PFC + 60% O_2_) versus control (*P* < 0.001, *n* ≥ 9). **(d)** Schematic illustration of the experimental design and treatment regimen in CT26 intradermal tumor model. BALB/c mice with 11-day established CT26 intradermal tumors were housed in 60% O_2_ chambers or maintained at 21% O_2_ with or without oxygenated Perflubron (10 mL/kg) 3 times per week until study completion. **(e)** Oxygenation agent therapy (PFC + 60% O_2_) induces the strongest tumor regression compared to control or either treatment alone. Evaluation of tumor growth kinetics and analysis is identical to (B). (*P<0.05, **P<0.005, (*) control versus PFC+60% O_2_; (#) control versus 60% O_2_. Data are presented as mean ±SEM, n ≥ 6). *Insert:* representative image of ulceration, necrosis and large holes observed in tumors from mice treated with oxygenation agent therapy. **(f)** Survival curves for CT26 tumor-bearing mice treated with oxygenation agent therapy (PFC + 60% O_2_) versus control (*P* < 0.001, n ≥ 6).

### Oxygenation agent therapy improves outcomes of adoptive cell therapy

The maximum therapeutic benefit of oxygenation agent therapy is likely to be achieved only when combined with other cancer immunotherapies, such as ACT ^43^. In this study, we suggest a feasible solution to one of the main impediments preventing therapeutic success of ACT in solid tumors – limited intratumoral infiltration/engraftment of T cells and suppression within the hostile TME. We, and others, have established that the hypoxic and adenosine rich TME inhibits infiltration of endogenous and adoptively transferred anti-tumor killer cells and downregulates effector functions and cytolytic activity ^7,8,16,28,29,59,60^. While many current cancer immunotherapy protocols assume that patients have sufficient anti-tumor immune cells, we suggest that only ACT may be capable of delivering high enough numbers to achieve clinical success ^60^. However, clinical outcomes of ACT in patients with solid tumors, even with chimeric antigen receptor (CAR) T cells, suggest that high numbers of transferred cells are insufficient due to inhibition by immunological and biochemical barriers in the TME.

To address this, we tested whether oxygenation agent therapy may reprogram the TME and enhance the therapeutic efficacy of adoptively transferred T cells. We utilized a well-established model of murine ACT in which mice bearing 11-day established pulmonary tumors were infused with 10×10^6^ culture-activated tumor-reactive T cells derived from tumor draining lymph nodes (TDLN) ^61,62^. One hour prior to T cell infusion, mice were treated with Perflubron, and again on days 12, 13, 14, 17 and 19 while housed in the hyperoxic chamber until the study completion on day 21. Fig. 5a,b and Supplemental Fig. 4 demonstrate that oxygenation agent therapy improves the efficacy of ACT since mice treated with this regimen in combination with transferred T cells exhibited significantly decreased numbers of metastatic nodules on the lungs compared to controls. While Perflubron and respiratory hyperoxia alone improve efficacy of transferred T cells, the combination therapy yielded the strongest tumor regression. Taken together, our data provide evidence that oxygenation agent therapy reprograms the TME via reversal of hypoxia to enhance the anti-tumor activity of adoptively transferred T cells. These preclinical data may provide justification for testing this strategy in cancer patients that are refractory to current cancer immunotherapies.

**Figure 5.**
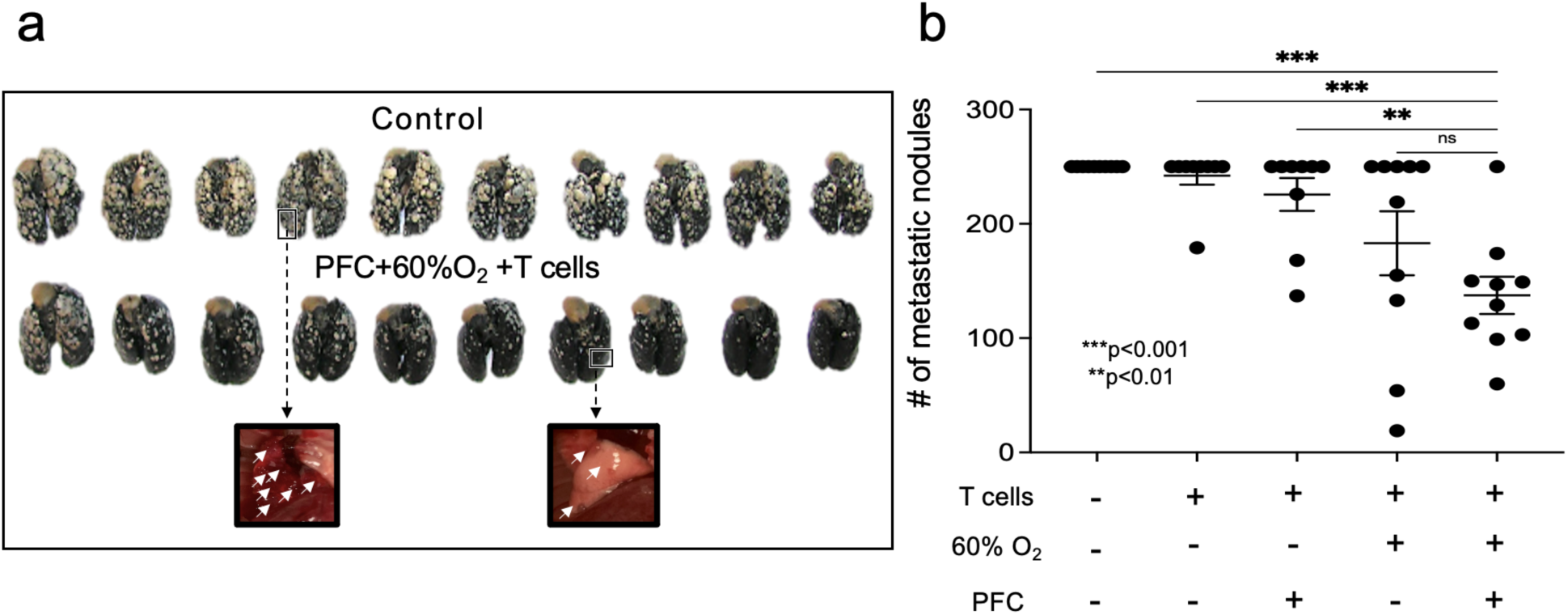
Oxygenation agent therapy improves efficacy of adoptive T cell therapy. Mice with 11-day established pulmonary metastases received adoptive cell transfer (ACT) of 10×10^6^ T cells. One day prior to ACT, mice underwent lymphodepletion with 100 mg/kg i.p. cyclophosphamide to mimic clinical protocols. Several hours prior to ACT, mice received intravenous administration of 10 ml/kg of PFC followed by five additional doses on day 12, 13, 14, 17 and 19. On day 11 (same day as ACT), mice were placed in 60% O_2_ chamber to maximize PFC O_2_ transport or maintained at 21% O_2_ as control until assay completion on day 21. After termination of the study, mice were sacrificed, and lung tumors enumerated by counterstaining with India ink. **(a)** Image of tumor-bearing lungs from control versus PFC+60% O_2_ +ACT (white arrows indicate metastatic nodules on the lungs prior to counterstaining). **(b)** Quantification of the number of metastatic nodules. Lungs with > 250 tumors were denoted as such since this is the maximum number that can be counted reliably. *P*-values are calculated using one-way ANOVA with post Tukey HSD (*P<0.05, **P<0.005, ***P<0.0005, ****P<0.0001. Data are presented as mean ± SEM, *n* ≥ 9).

### Oxygenation agents in cancer patients unable to receive respiratory hyperoxia

Our data suggest that the combination of oxygen-carrying Perflubron with respiratory hyperoxia is more effective in inducing tumor regression than either strategy alone. However, there may be clinical situations where the use of respiratory hyperoxia may be limited, especially in patients who would not benefit from systemic exacerbation of the immune system, such as those with lung injuries, ongoing inflammation, or active autoimmunity. In addition, some patients may elect not to remain on respiratory hyperoxia continuously following treatment.

Therefore, we asked whether Perflubron administered without respiratory hyperoxia may also improve the efficacy of adoptive T cell therapy. Mice with 11-day established MCA205 fibrosarcoma pulmonary tumors received 10×10^6^ T cells derived from tumor draining lymph nodes. One hour prior to ACT, mice were treated with Perflubron, and again on days 13, 15 and 17. On day 19, mice were sacrificed, and pulmonary tumors were enumerated. Fig. 6a,b and Supplemental Fig. 6 demonstrate that perflubron administration alone can improve the therapeutic efficacy of adoptive T cell therapy since mice receiving Perflubron during ACT demonstrated significantly improved tumor regression compared to control animals. Interestingly, Fig. 6A (*left*) and Supplemental Fig. 5 indicate that Perflubron alone (without ACT or respiratory hyperoxia) is able to reduce lung tumor burden, although most lungs had greater than 250 metastases which is the highest that may be counted reliably. Taken together, data from Fig. 5 and 6 suggest that treating patients with oxygenation agent therapy prior and during ACT can reprogram the TME by reversing hypoxia-adenosinergic suppression and improve therapeutic responses.

**Figure 6.**
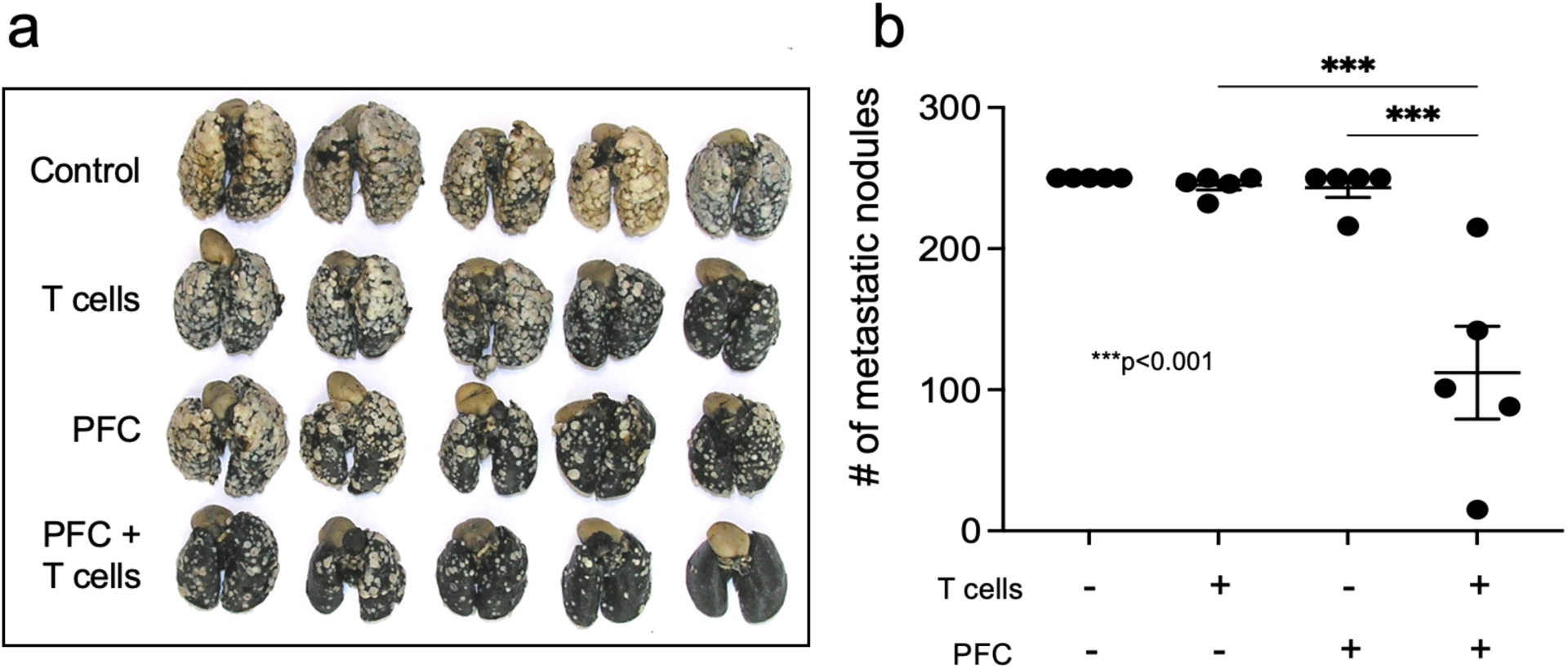
Perflubron administration alone is capable of improving efficacy of adoptive T cell therapy. Mice with 11-day established pulmonary metastases received adoptive cell transfer of 10×10^6^ T cells. One day prior to ACT, mice underwent lymphodepletion with 100 mg/kg i.p. cyclophosphamide. Several hours prior to ACT, mice received intravenous dose of 10 ml/kg of PFC followed by three additional doses on days 13, 15 and 17. At the completion of the study (day 19), mice were sacrificed, and metastatic nodules were enumerated by counterstaining with India ink. **(a)** Image of tumor bearing lungs from each treatment group. **(b)** Quantification of the number of metastatic nodules. *P*-values are calculated using one-way ANOVA with post Tukey HSD (*P<0.05, **P<0.005, ***P<0.0005, ****P<0.0001. Data are presented as mean ± SEM, n = 5)

## Discussion

These preclinical studies provide evidence to justify the repurposing of available, safe, and well-tolerated blood substitutes to address the medical need in improving existing cancer therapies. ACT using CAR T cells and TCR T cells may represent the most powerful immunotherapeutic approach to date since it ensures the presence of sufficient numbers of tumor-reactive immune cells. However, the use of ACT in the clinic has been challenging, particularly in the treatment of solid tumors that are protected by immunological and biochemical negative regulators. While elimination of immunological barriers by PD-1/CTLA-4 blockade have yielded improved responses to some cancers, immune cells are still inhibited by the biochemical barriers derived from oxygen-poor and extracellular adenosine-enriched TMEs. Subsequently, the absence of anti-tumor killer cells in regions of tumor hypoxia weakens the therapeutic efficacy of immune checkpoint blockade (ICB) and ACT providing justification to target hypoxia to reprogram the TME and improve the success of cancer immunotherapies.

Here, we introduce the conceptually novel approach of eliminating the powerful immunosuppressive barrier governed by hypoxia/HIF-α and adenosine → A2AR → cAMP-mediated signaling and immunosuppressive transcription by using oxygenation agents. The novelty of this approach is in utilizing blood substitutes to reprogram the TME by increasing oxygen concentration and weakening the upstream stage of the hypoxia-adenosinergic axis (Fig. 1a). The advantage of blood substitutes exists in their significantly smaller size compared to erythrocytes, providing the ability to carry oxygen farther away from the blood vessels and deeper within the disorganized and aberrant tumor microvasculature. This dramatically improves the ability to oxygenate tumors that are in different anatomical locations when compared to more oxygen privileged organs, such as the lungs ^28^.

These data provide proof of principle for the use of a class of oxygenation agents (perfluorocarbons) to improve cancer immunotherapy by showing that Perflubron: i) carries and delivers oxygen into tumors, ii) eliminates hypoxia in the TME, iii) improves tumor regression and survival and iv) enhances therapeutic efficacy of adoptive T cell transfer. Additionally, we provide here a methodological approach to increase the amount of dissolved oxygen in PFCs prior to *in vivo* infusion by saturation with 100% O_2_, and post infusion by exposure to higher fractions of inspired O_2_ achieved by respiratory hyperoxia (60% O_2_).

Finally, our studies further elucidate the relationship between the hypoxic TME and infiltration of tumor reactive T cells and NK cells. Interestingly, we found a stronger negative correlation between infiltration of NK cells and hypoxic levels when compared to that of CD8 T cells. Our future work aims to clarify the extent to which NK cells may be more sensitive to immunosuppressive hypoxic signaling, which is in accordance with previous studies demonstrating a high susceptibility of NK cells to A2AR-mediated inhibition ^28,29,57,58^. Indeed, the inhibition of T cells and NK cells by hypoxia/HIF-1α and adenosine → A2AR → cAMP-mediated transcription is supported by our previous work ^24,25,27-31,43^. These studies were the first to establish that hypoxic reversal in the TME using respiratory hyperoxia reduces the transcription of immunosuppressive genes associated with HRE and CRE activity, resulting in a reduction in extracellular adenosine, decreased expression of adenosine receptors and adenosine-generating ectoenzymes CD39/CD73, and decreased immunosuppressive molecules (e.g. TGF-β) and reduced suppressor cells (e.g. T regs) ^28,29^. Such observations along with the current understanding of the absence of tumor reactive T cells and NK cells from hypoxic regions in the TME provide rationale for combining oxygenation agent therapy with cancer immunotherapy protocols.

Data from this study, together with our previous work ^7,27-29^ support the following immunotherapeutic protocol where patients with refractory solid tumors may be treated with i) adoptively transferred T cells (e.g. TCR or CAR T cells), ii) monoclonal antibody to inhibit immunological negative regulators, iii) drugs that eliminate extracellular adenosine-A2AR-cAMP mediated signaling, and iv) oxygenation agents that eliminate or weaken the hypoxia-HIF1α signaling that triggers downstream HRE transcriptional programming and immunosuppressive adenosine-cAMP axis. In accordance with such a protocol are recent updates from Phase I clinical trials (NCT03098160) in which 66.7% of patients receiving combination of evofosfamide (hypoxia-activated prodrug) and ipilimumab achieved stable disease and 16.7% achieved partial response^50^. Additionally, patients with pre-existing immune gene signatures have been predicted to respond to therapy which further highlights the necessity of ACT for success of cancer immunotherapies ^23,50,63,64^.

Our strategy to target tumor hypoxia to improve cancer immunotherapy is also supported by pioneering work by Semenza’s group using pharmacological agents to target HIFs ^65,66^. Recently, in a model of murine hepatocellular carcinoma, characterized by high intratumoral hypoxia and poor immunotherapeutic responses, the addition of the HIF inhibitor 32-134D to anti-PD1 therapy increased the rate of tumor eradication from 25% to 67% ^66^. Similar to our observations reported here and in our previous work ^28,29^, inhibition of HRE-driven transcription led to a reduction in suppressor cells (tumor-associated macrophages and myeloid-derived suppressor cells) and an increase in CD8 T cells and NK cells ^66^. Our future work will determine whether oxygenation agent therapy may improve responses to immune checkpoint blockade.

Importantly, oxygenation via PFCs may benefit from higher fraction of inspired oxygen to maximize O_2_ transport capacity ^54,67,68^. This study identifies a unique treatment modality combining PFC with respiratory hyperoxia (60% O_2_, which previously demonstrated anti-tumor efficacy by weakening hypoxia-adenosinergic immune-suppression) to improve anti-tumor immune responses against solid tumors ^28,29^. We also show for the first time that an oxygen-delivering nanoemulsion can improve ACT and may also be a powerful method to improve other forms of cancer immunotherapy ^43^. Extension from this work suggests that treating patients with oxygenation agents prior to ACT may reprogram the TME before arrival of transferred T cells or CAR T cells to improve infiltration, activation, and effector responses. Thus, oxygen delivery via PFC may offer a safe and effective strategy to target hypoxia-driven immunosuppressive barriers in the TME. Importantly, this treatment option can be readily available as previous investigation into blood substitutes for several other clinical indications were shown to be safe and well tolerated in over 3,000 patients ^54^.

## Materials and Methods

### Animals

All animal experiments were approved by the Institutional Animal Care and Use Committee (IACUC) guidelines of Northeastern University. Female C57BL/6 or BALB/c wild type (WT) mice (8-12 weeks old) were purchased from the Jackson Laboratory. Female mice were used for the tumor immunology assays described since they exhibit more consistent immune responses in these models. Animals were housed in a specific-pathogen free environment in a 12h-12h (light-dark) conditions in approximately 24°C according to the guidelines from National Institute of Health (NIH).

### Murine tumor models

Two different tumor models were employed in these studies: i) the weakly immunogenic MCA205 fibrosarcoma cell line of B6 origin and ii) the more immunogenic (“hot tumor”) CT26 colon carcinoma purchased from American Type Culture Collection (ATCC). Both cell lines were administered intradermally (i.d.) at doses described in each study and mice were randomized following tumor administration. To provide models of different anatomical location, MCA205 fibrosarcoma was also administered intravenously for experimental pulmonary metastasis. Both cell lines were maintained in complete media (CM) containing RPMI-1640 (Bio Whittaker) supplemented with 10% heat-inactivated fetal bovine serum (FBS), 100U/mL penicillin, 100µg/mL streptomycin (Gibco), 0.1mM non-essential ammino acids (Gibco), 1µM sodium pyruvate (Gibco), 2mM L-glutamine (Gibco, Fisher Scientific), 5×10^-5^ M 2-mercaptoethanol (Gibco) and 50µg/mL Gentamicin (Lonza, Fisher Scientific). Cells were cultured at 21%O_2,_ 5%CO_2_ and 37°C in Hera Cell Vios 160i incubators (ThermoFisher). In vivo inoculation occurred between 4-6 passages.

### Oxygenation agent therapy

#### Respiratory hyperoxia

For studies employing respiratory hyperoxia, mice were housed in chambers with well-controlled gas composition to mimic clinical protocols as done previously ^28,29^. Hyperoxia in the chamber is maintained with self-contained oxygen generators (AirSep) to ensure consistent oxygen levels and monitored continuously using oxygen Alpha Omega Instruments oxygen sensors. To avoid hypercapnic acidosis, traditional cage lids were replaced with aerated wire lid and SodaSorb (Medline) was added to absorb any excess carbon dioxide inside the chamber as done previously ^28,29^ ^69-71^. Fractional concentrations of O_2_ and CO_2_ inside the chambers was determined previously by pulling a sample from the chamber at a rate of 100 ml min−1 using a Sable Systems Model SS3 sample pump (Sable Systems) as described ^28,29^.

#### Perflubron preparation for in vivo administration

Dissolved oxygen is approximately 21% O_2_ in “off the shelf” Perflubron since the amount of dissolved O_2_ in perfluorocarbon-based oxygen carriers is directly proportional to the partial pressure of O_2_ ^54^ (Fig. 3). To maximize oxygen-carrying capacity prior to in vivo administration, Perflubron was infused with 100% O_2_ for seven minutes and then sealed in an air-tight container with a fitted rubber stopper. To improve injectability, Perflubron was diluted 1:1 in 1 x Hank’s Balanced Salt Solution (Hy Clone, Fisher Scientific) prior to saturation with 100% O_2_. To minimize reduction of oxygen in the sealed container, Perflubron was withdrawn with a 27-gauge needle for intravenous (i.v.) administration (200 μl) into each mouse and then immediately re-sealed with parafilm.

### Evaluation of intratumoral hypoxia and the tumor microenvironment

C57BL/6 female mice were injected intradermally (i.d.) with 0.1 × 10^6^ MCA205 fibrosarcoma cells and then randomized. On day 7 after tumor inoculation, mice were placed into the hyperoxic chambers (60% O_2_) for 72 h or kept at atmospheric oxygen levels (21% O_2_). Tumor bearing mice receiving oxygenation agent therapy received i.v. Perflubron (15mL/kg) on days 7, 8 and 9 of tumor growth. One hour before sacrifice, mice were injected intraperitoneally (i.p.) with 80mg/kg of Hypoxyprobe^™^-1 (pimonidazole HCl) (Hypoxyprobe, Inc., MA) dissolved in HBSS. Pimonidazole creates thiol protein adducts in tissues with pO_2_ ≤ 10mmHg. On day 9, mice were sacrificed, and tumors were evaluated for intratumoral hypoxia by immunofluorescence staining of frozen tumor sections and flow cytometry.

#### Immunofluorescence analysis of the TME

For immunofluorescence analysis, tumors were frozen in Tissue-Tek Optimal Cutting Temperature (O.C.T) compound (Sakura, Fisher Scientific) and 5 µm thick sections were mounted on polypropylene coated glass slides (Thermo Fisher) from 10 to 20 different cutting surfaces. Slides were air dried for 45 minutes and fixed in 1:1 mixture of acetone and methanol for 10 min at -20°C and stored at -20°C. To detect levels of hypoxia and endogenous immune cell infiltration, slides were first labeled with Fc block (BD Pharmingen) at 1:100 in staining buffer (1x PBS, 0.5% bovine serum albumin, 0.1% Tween 20) for 10 min at RT in the dark followed by staining with FITC anti-Hypoxyprobe mAb (Hypoxyprobe^TM^), PE anti-CD8 (clone 53-6.7, Biolegend) and APC anti-NK1.1 (clone PK136, Biolegend) mAbs in concentration 1:100 for 3 hours. Slides were washed and stained for 5min with 4′,6-diamidino-2-phenylindole (DAPI, ThermoFisher) for detection of nuclei and mounted with Fluoromount-G (SouthernBiotech). Fluorescently labeled slides were imaged using IX83 microscope (Olympus) and analyzed using CellSens Software (Olympus).

#### Flow cytometry analysis of the TME

Flow cytometric analysis performed on isolated tumors that were cut into small pieces (2-4 mm) and homogenized using Murine Tumor Dissociation kit (Miltenyi) and gentleMACS Dissociator (Miltenyi) following manufacturers protocol for homogenization of soft/medium tumors. After homogenization, cells were resuspended in 40% Percoll^TM^ prepared by dilution of 100% Percoll (Cytiva) in CM pre-warmed to RT and centrifuged for 10 min at 3,000 x g. Lymphocytes and tumor cells were pelleted and debris was separated at the top and removed by aspiration. To remove red blood cells (RBCs), cell suspension was mixed with 3 mL of ACK Lysing Buffer (Gibco). Remaining cells were stained and acquired using a FACSCalibur Cytek DxP 8 and analyzed with FlowJo. Cells were first washed with staining buffer (1xPBS, 1%BSA, 1% penicillin streptomycin) and centrifuged at 3000 RPM for 3 min to obtain a cell pellet at the bottom of the tube. Wash was aspirated leaving the pelleted cells in approximately 40uL of staining buffer. Samples were first stained with Fc Block cocktail (BD Pharmingen) containing 9.5µL staining buffer and 0.5µL Fc Block (BD Pharmingen) for 10min at 4°C. Without washing, 20µL of surface mAb cocktail was directly added to the cell suspension. Cells were stained for surface markers for 30 min at 4°C. For detection of hypoxia, samples were fixed and permeabilized using eBioscience^TM^ FoxP3/Transcription Factor Staining Buffer Set (Invitrogen) following manufacturer’s protocol. Intracellular staining was performed with FITC anti-Hypoxyprobe (Hypoxyprobe, Inc., MA) mAb for 1h at RT in permeabilization buffer. Following staining, cells were washed twice and resuspended in staining buffer, passed through 70 µM filter (Corning) and analyzed on FACSCalibur Cytek DxP 8. Data were analyzed using FlowJo software.

### Real-time Perflubron-mediated elevation of intratumoral oxygen levels

C57BL/6 female mice were inoculated i.d. with 5×10^5^ MCA205 fibrosarcoma cells and randomized. When tumors reached approximately 6mm × 7mm, tumor bearing mice were anesthetized and the tumor was probed using Presens Oxygen Monitoring System and Oxygen Sensors (PreSens). After identification of a hypoxic region in the TME (O_2_ < 1%) and stabilization of the readout (∼12 min), mice received i.v. administration of oxygenated Perflubron (15 mL/kg). Oxygen concentration was recorded for ∼4 h (229.5 min) post infusion. Experiment was repeated three times.

### Tumor growth kinetics and survival studies

Female C57BL/6 mice were inoculated i.d. with 0.1×10^6^ MCA205 fibrosarcoma cells resuspended in 100µL HBSS and randomized. Oxygenation agent therapy was initiated on day 9. For CT26 colon carcinoma model, 0.75×10^5^ tumor cells were injected intradermally into BALB/c WT and oxygenation agent therapy was initiated on day 11. For both in vivo tumor models, oxygenation agent therapy consisted of housing animals in hyperoxic chambers (60% O_2_) and 3 injections/week of 10mL/kg of Perflubron (i.v.) until study completion. Tumors growth kinetics were measured in a blinded manner 3x per week using Vernier calipers and volume was calculated using the following formula: 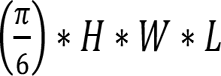.

### Adoptive Cell Transfer (ACT)

#### Preparation of culture activated tumor-draining lymph node (TDLN) T cells for ACT

Female C57BL/6 donor mice were injected subcutaneously into both flanks with 1×10^6^ MCA205 fibrosarcoma cells resuspended in 100µL HBSS. After twelve days, inguinal tumor draining lymph nodes (TDLN) were isolated and processed to be activated with plate bound anti-CD3 as described previously ^28^. One day prior, wells were coated with Protein A from *Staphylococcus aureus* (Sigma). After 48h, cells were collected and expanded in culture at 0.35 × 10^6^ per ml and supplemented with 100IU IL-2 for 96 hours in gas permeable MACS^R^ GMP Cell Differentiation Bags (Miltenyi). After expansion, cells were collected, washed, and resuspended in HBSS. Therapeutic efficacy of transferred T effector cells was assessed in the treatment of 11-day established MCA205 pulmonary tumors by intravenous injection of 10×10^6^ culture-activated T cells to each mouse. Recipient mice were injected i.v. with 2×10^5^ MCA205 tumor cells at day 0 for lung tumor establishment. Recipient mice were also pretreated with cyclophosphamide (100 mg/kg i.p.) 1 day before infusion of T cells. Cyclophosphamide treatment is routinely used to improve the therapeutic efficacy of adoptively transferred T cells and was also administered to untreated tumor-bearing control mice ^61,62^.

#### ACT and Perflubron in the treatment of pulmonary metastases

Recipient mice were injected i.v. with 2×10^5^ MCA205 tumor cells for lung tumor establishment and randomized. One day prior to adoptive T cell transfer, tumor-bearing recipient mice received lymphodepletion with 100mg/kg of cyclophosphamide. On day 11 of tumor growth, mice received i.v. infusion of 10×10^6^ T cells. Several hours prior to ACT, mice were given oxygenated Perflubron i.v. (10mL/kg). Mice received additional doses of Perflubron (10mL/kg) on day 13,15 and 17 until the study completion on day 19. Mice were sacrificed, tumor-bearing lungs were counterstained with India ink and tumors were enumerated in a blinded manner. Lungs with more than 250 nodules were assigned >250 as this is the maximum number that can be counted reliably.

#### ACT and oxygenation agent therapy in the treatment of pulmonary metastases

Study design is identical to above except on day 11 mice were housed in hyperoxic chambers [respiratory hyperoxia (60% O_2_)] or under atmospheric oxygen levels (21% O_2_) until completion of the study on day 21. During this time, mice were treated with or without Perflubron (10ml/kg) on days 11, 13, 15, 17, and 19. On day 21, mice were sacrificed, tumor-bearing lungs were counterstained with India ink and tumors were enumerated in a blinded manner. Lungs with more than 250 nodules were assigned >250 as this is the maximum number that can be counted reliably.

### Statistics

The significance of differences in the numbers of infiltrating CD8 and NK cells into hypoxic vs non-hypoxic regions was evaluated using the student’s t test (two-sided). Differences between treatment groups in immunofluorescence staining of frozen tumor sections and flow cytometric assays was evaluated using one-way ANOVA. For studies of tumor growth kinetics, differences between groups were evaluated using two-way ANOVA (mixed-effects analysis). All survival studies were analyzed using Log-rank test. Significance of differences of lung tumors between groups was evaluated using one-way ANOVA. All P values are listed within the figures or figure legends.

### Study Approval

All animal studies were approved by the Institutional Animal Care and Use Committee (IACUC) guidelines of the Division of Laboratory and Animal Medicine at Northeastern University

## Data Availability

The data generated in this study are available upon request from the corresponding author.

## List of Supplementary Materials

Fig S1 to S5 are presented along with figure legends in Supplementary Materials

## Acknowledgements

Perflubron was shared by Dr. Bruce Spiess via material transfer agreement from the University of Florida, Gainesville.

## Author contributions

K.V.H performed experiments, participated in study design and analysis, and wrote the manuscript together with S.M.H. S.M.H. led the study from conception to design of experiments, provided troubleshooting and analysis of data. M.V.S provided institutional memory of anti-hypoxia-adenosinergic approach and infrastructure at the New England Inflammation and Tissue Protection Institute. M.V.S. also provided constructive criticism and participated in manuscript preparation. B.S. provided institutional knowledge of blood substitutes and oxygenation agents, provided Perflubron nanoemulsions used in the studies, and edited the manuscript. M.M., A.A, N.R, C.B., M.S., K.W., and L.R. assisted in execution of experiments and participated in discussion and analysis of results.

