## Supplementary Figures for "Oxygen carrying nanoemulsions and respiratory hyperoxia eliminate tumor hypoxia-induced suppression and improve cancer immunotherapy"

### Supplementary Materials

#### Supplemental Figures

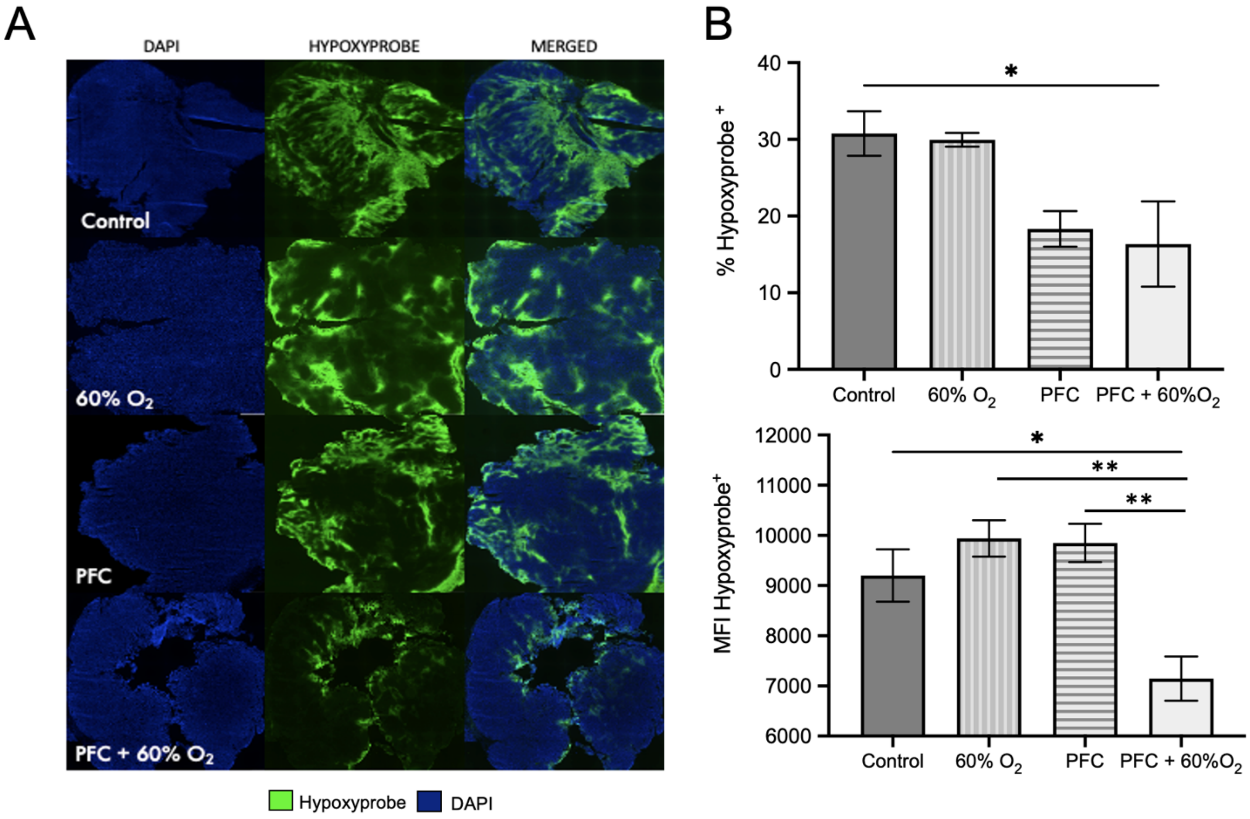


**Supplemental Fig. 1. Oxygenation agent therapy reduces intratumoral hypoxia.** Mice with 8-day established MCA205 intradermal tumors were treated with or without i.v. administration of 15 ml/kg oxygenated Perflubron (PFC) for 72 h with continuous breathing of 60% $O_2$ (respiratory hyperoxia) or 21% $O_2$ (normoxia).

**(A)** Representative images of all experimental groups showing the decrease of intratumoral hypoxia after 72 h treatment respiratory hyperoxia (60% $O_2$), Perflubron (PFC), and oxygenation agent therapy (PFC + 60% $O_2$) compared to control.

**(B)** Flow cytometric analyses demonstrating the percent of Hypoxyprobe positive cells (top) as well as mean fluorescent intensity (MFI) of Hypoxyprobe positive cells (bottom). While both PFC and PFC + 60% $O_2$ groups showed a decrease in the percentage of hypoxic cells, only mice treated with PFC + 60% $O_2$ exhibited near complete reduction of hypoxia and reduced hypoxyprobe MFI. Significance calculated using one-way ANOVA with Tukey post-test for multiple comparisons (*P<0.05).

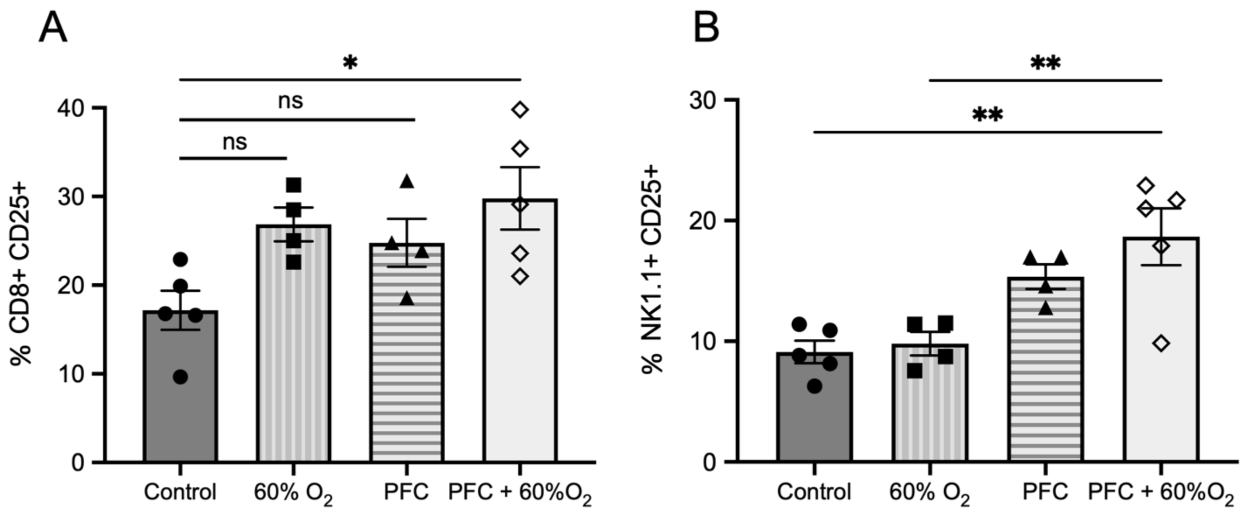


**Supplemental Fig. 2. Oxygenation agent therapy reprograms the tumor microenvironment toward anti-tumor immunity.** C57BL/6 mice with 8-day established MCA205 intradermal tumors received daily administration (i.v., 15 ml/kg) of oxygenated Perflubron for 72 h while breathing 60% $O_2$ (respiratory hyperoxia) or 21% $O_2$ (normoxia). Mice were injected intraperitoneally with Hypoxyprobe-1™ (80mg/kg) one hour before sacrifice to determine levels of intra-tumoral hypoxia. After sacrifice, tumors were isolated divided for analysis via immunofluorescence or flow cytometry. The percent CD8+CD25+ **(A)** and NK+CD25+ **(B)** cells from tumor digests of all experimental groups as measured by flow cytometry (*n* = 5 for Control, PFC + 60% $O_2$ , *n* = 4 for PFC, 60% $O_2$). *P*-values are calculated using one-way ANOVA with Tukey post-test for multiple comparisons (*P<0.05). Data are presented as mean ±SEM.

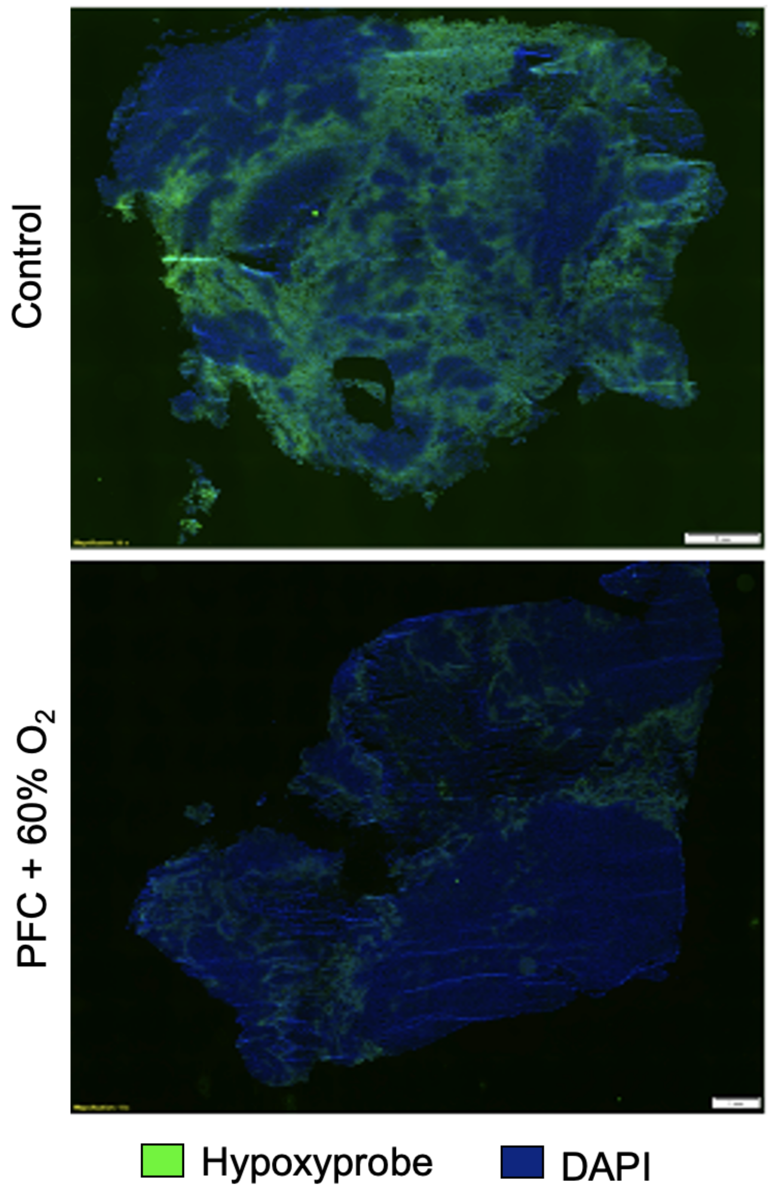


**Supplemental Fig. 3. Oxygenation agent therapy induces potent tumor regression and eliminates hypoxia in remaining tumors.** C57BL/6 were injected intradermally with MCA205 fibrosarcoma and treated with or without Perflubron (PFC) and respiratory hyperoxia (60% $O_2$)**.** Oxygenation agent therapy (PFC + 60% $O_2$) induces the strongest tumor regression compared to control or either treatment alone (Fig. 6B). At study endpoint mice were injected intraperitoneally with 80mg/kg of Hypoxyprobe to evaluate hypoxic regions in treated and untreated tumors. Isolated tumors were frozen in OCT and cut to 5 uM thick sections and labelled with anti-Hypoxyprobe antibody. Representative images of tumors from control mice (top) and oxygenation agent therapy treated mice (bottom) are shown (10x magnification, scale bar represents 1 mm). Remaining tumors from PFC + 60% $O_2$ treated mice exhibited dramatically less hypoxic staining at study endpoint.

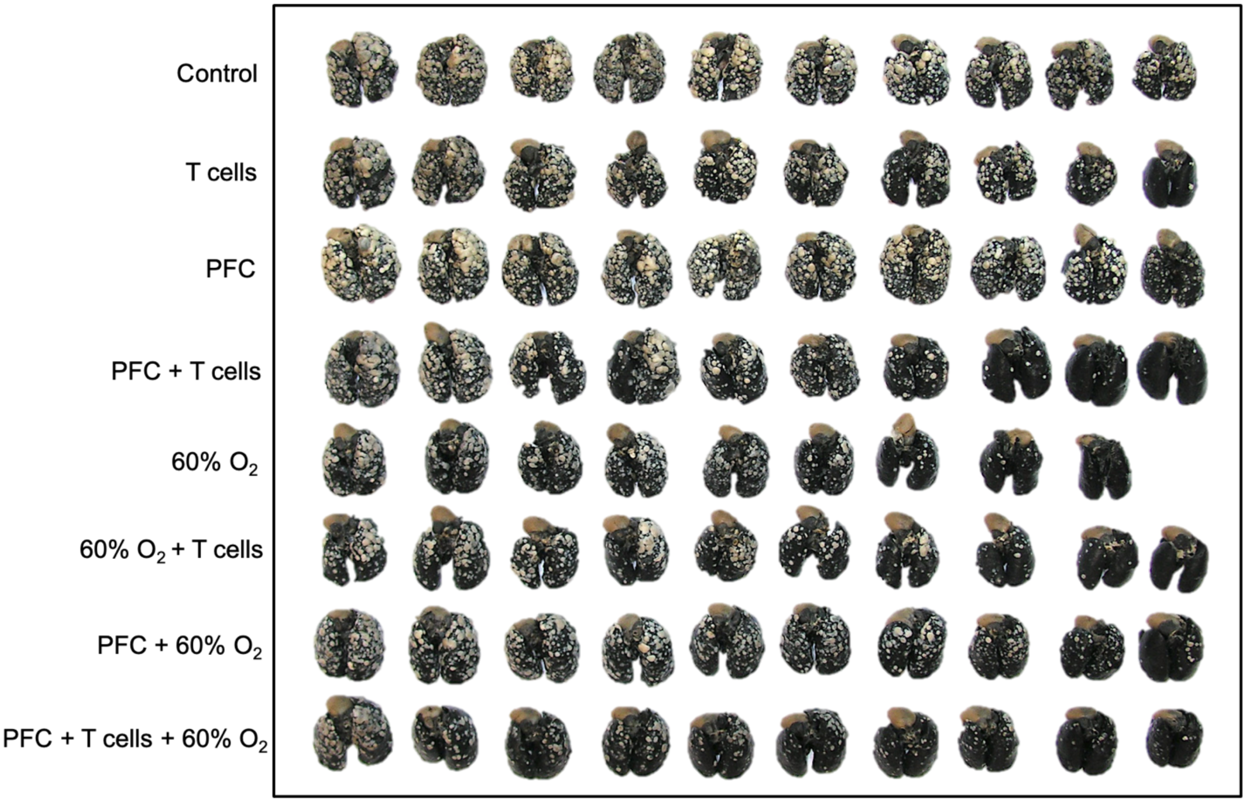


**Supplemental Fig. 4. Oxygenation agent therapy improves efficacy of adoptive cell transfer.** Mice with 11-day established pulmonary metastasis received adoptive cell transfer (ACT) of $10 \times 10^6$ T cells. One day prior to ACT, mice underwent lymphodepletion with 100 mg/kg i.p. cyclophosphamide to mimic clinical protocols. Several hours prior to ACT, mice received intravenous administration of 10 ml/kg of PFC followed by five additional doses on day 12, 13, 14, 17 and 19. On day 11 (same day as ACT), mice were placed in 60% $O_2$ chamber to maximize PFC $O_2$ transport or maintained at 21% $O_2$ as control until assay completion on day 21. After termination of the study, mice were sacrificed, and counterstained with India ink. Representative images of tumor bearing lungs from all experimental groups are shown.

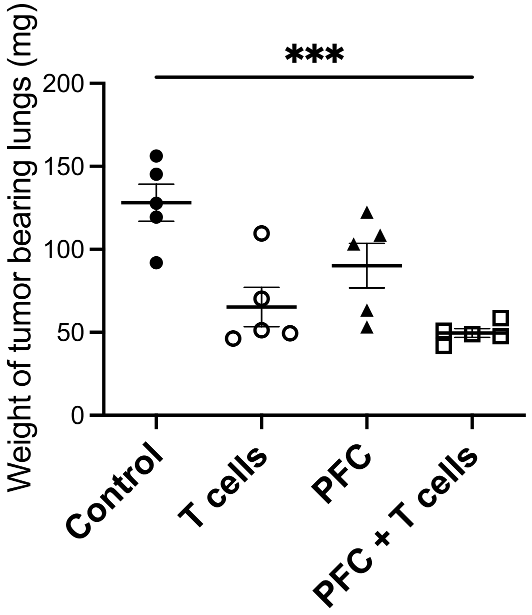


**Supplemental Fig. 5. Perflubron administration alone is capable of improving efficacy of adoptive cell transfer.** Mice with 11 days established pulmonary metastases received adoptive cell transfer of $10x10^6$ T cells. One day prior to ACT, mice underwent lymphodepletion with 100 mg/kg i.p. cyclophosphamide. Several hours prior to ACT, mice received intravenous dose of 10 ml/kg of PFC followed by three additional doses on days 13, 15 and 17. At the completion of the study (day 19), mice were sacrificed, and lung tumors enumerated. Following enumeration of surface tumors, lungs were dried and weighed to assess total lung tumor burden. *P*-values are calculated using one-way ANOVA with post Tukey HSD (***$P<0.0005$, data are presented as mean ± SEM, $n \geq 9$).
